# The sensitivity to Hsp90 inhibitors of both normal and oncogenically transformed cells is determined by the equilibrium between cellular quiescence and activity

**DOI:** 10.1101/472290

**Authors:** Pablo C. Echeverria, Kaushik Bhattacharya, Abhinav Joshi, Tai Wang, Didier Picard

## Abstract

The molecular chaperone Hsp90 is an essential and highly abundant central node in the interactome of eukaryotic cells. Many of its large number of client proteins are relevant to cancer. A hallmark of Hsp90-dependent proteins is that their accumulation is compromised by Hsp90 inhibitors. Combined with the anecdotal observation that cancer cells may be more sensitive to Hsp90 inhibitors, this has led to clinical trials aiming to develop Hsp90 inhibitors as anti-cancer agents. However, the sensitivity to Hsp90 inhibitors has not been studied in rigorously matched normal versus cancer cells, and despite the discovery of important regulators of Hsp90 activity and inhibitor sensitivity, it has remained unclear, why cancer cells might be more sensitive. To revisit this issue more systematically, we have generated an isogenic pair of normal and oncogenically transformed NIH-3T3 cell lines. Our proteomic analysis of the impact of three chemically different Hsp90 inhibitors shows that these affect a substantial portion of the oncogenic program and that indeed, transformed cells are hypersensitive. Targeting the oncogenic signaling pathway reverses the hypersensitivity, and so do inhibitors of DNA replication, cell growth, translation and energy metabolism. Conversely, stimulating normal cells with growth factors or challenging their proteostasis by overexpressing an aggregation-prone sensitizes them to Hsp90 inhibitors. Thus, the differential sensitivity to Hsp90 inhibitors may not stem from any particular intrinsic difference between normal and cancer cells, but rather from a shift in the balance between cellular quiescence and activity.

## Introduction

From its discovery almost four decades ago, the molecular chaperone heat-shock protein 90 (Hsp90) was considered a protein assisting oncogenic processes [1,2]. An extensive literature establishes the essential role of Hsp90 in development and differentiation at both the cellular and organismic levels, in health and disease, in hosts and pathogens. A complete overview of facts and literature on Hsp90 can be found here: https://www.picard.ch/downloads/Hsp90facts.pdf. Whenever a new cellular stage, process, transcriptional program or regulatory state is engaged, Hsp90 is present to assist it. Hsp90 is at the center of the cellular proteome acting as a major hub sustaining a vast number of proteins and protein-protein interaction networks that maintain cellular homeostasis and function [3-5]. A relevant example of that is the fact that Hsp90 allows, supports and maintains neoplastic transformation; qualitative and quantitative changes of the protein network of cancer cells appears to make them more dependent on the Hsp90 molecular chaperone machine [6-8].

Hsp90 functions as a dimer and requires complex ATPase-associated conformational changes regulated by a large spectrum of co-chaperones to process its substrates, also referred to as its clientele [9]. Due to unique features of the N-terminal ATP binding pocket of Hsp90, specific competitive inhibitors of Hsp90 have been developed [10,11]. Intriguingly, cancer cells were found to be more sensitive to Hsp90 inhibitors than normal cells, conceivably reflecting their increased dependency on Hsp90 chaperone activity; however, although a number of molecular mechanisms have been proposed to explain this difference, the precise reasons have remained controversial and incompletely characterized [12-22]. The difference in sensitivity suggested a therapeutic window and prompted high hopes for the treatment of cancer and led to a large number of clinical trials. 20 years later, there is still no Hsp90 inhibitor in the clinic and only few ongoing clinical trials left [23,16,24]. There may be many reasons for this frustrating outcome, but there is a general sense that Hsp90 inhibitors were not properly evaluated before reaching the clinic, that good biomarkers of successful Hsp90 inhibition were missing, and that patients should have been stratified more carefully [25].

In revisiting the published cellular models for comparing Hsp90 inhibitor sensitivities, it occurred to us that there is a serious problem of comparing “apples and oranges”. We therefore decided to adopt an old model of oncogenic transformation [26]. Mouse NIH-3T3 fibroblast cells can be considered normal cells except for the fact that they are immortal. Upon transformation by oncogenes, they acquire many hallmarks of cancer cells, including the ability to form tumors in immunocompromised mice. This allowed us to generate an isogenic pair of normal (N) and Ras-transformed NIH-3T3 fibroblast cells (R). Hence, the latter differs from the former by a single gene. We extensively characterized this pair of cell lines with regards to the proteome changes, sensitivity to Hsp90 inhibitors, and impact of these inhibitors. These analyses showed them to constitute a powerful model to begin to understand the molecular mechanisms underlying the therapeutically highly relevant differential sensitivity of normal and cancer cells to Hsp90 inhibitors.

## Materials and methods

### Reagents

Antibodies used: GAPDH (HyTest Ltd., clone 6C5, cat. # 5G4), caspase-3 (Abcam, ab4051), Cdk6 (Abcam, ab3126), Ras (Stressgen, now Enzo Life Sciences; ADI-KAP-GP001), p-ERK (Santa Cruz, sc-7383), Hsp70 (Abcam, ab181606), p27Kip1 (Cell Signaling, #2552), Nr2f1 (ABGENT, #AP14218a), tubulin (Merck, CP06), Cdk4 (NeoMarkers, MS-616-P1), Raf-1 (Santa Cruz Biotechnology, sc-133), Akt (Cell Signaling, #9272). Transfection reagents: jetPRIME (Polyplus, #114-15), polyethylenimine (PEI) (Polysciences, # 621405). Hsp90 inhibitors: geldanamycin (LC Laboratories, G-4500), PU-H71 (Tocris, #3104), pochoxime A (kindly provided by Nicolas Wissinger, University of Geneva). Other inhibitors: AZ628 (Tocris, #4836), PD98059 (Tocris, #1213), cycloheximide (Sigma, C7698), rapamycin (LC Laboratories, R-5000), aphidicolin (ACROS, BP615-1), hydroxyurea (fluorochem, #043351). All compounds were dissolved in DMSO. Growth factors: insulin (Sigma, I6634), recombinant human epidermal growth factor (EGF) (Lonza, CC-4017). Metabolic assay reagents: Oligomycin (SIGMA, 7535), carbonyl cyanide-p-trifluoromethoxyphenylhydrazone (FCCP) (SIGMA, C2920) and rotenone (SIGMA, R8875) with antimycin (SIGMA, A8674).

### Cells, cell culture, plasmids and transfection

Cells were routinely cultured in Dulbecco’s modified Eagle’s medium (DMEM) supplemented with 10% heat-inactivated fetal bovine serum and penicillin/streptomycin. R cell clones were obtained by transfection of N cells in 6-well plates using polyethyleneimine (PEI) at a DNA-to-PEI ratio of 1:2 with the plasmid pBW1631, which contains a genomic fragment of the *Ha-c-Ras* gene with a point mutation encoding the alteration G12V from T24 bladder carcinoma (a gift from Frank McCormick) [27]. 48 hrs after transfection, cells were trypsinized and plated in a 100 mm plate. The culture medium was changed every 2 days. 15 days after transfection, transformed foci were apparent and could be selected and established as stable clones. For proteostasis distress assays, N cells were transfected with plasmids pEGFP-Q74 (for expression of the aggregation-prone form) and pEGFP-Q23 (for expression of the non-aggregating form), which were a gift from David Rubinsztein (Addgene plasmids # 40262 and # 40261) [28]. N cells were transfected using the jetPRIME transfection reagent following the manufacturer’s instructions.

### Metabolic assays

Glycolysis was monitored in real time using a seahorse XF analyzer (XF24, Agilent). 4 x 10^4^ Rc and N cells were cultured overnight in custom XF24 microplates either with complete DMEM with all major carbon sources (glucose, pyruvate, glutamine) or DMEM with only glucose as the carbon source. Before the measurements, all cells were washed & incubated in unbuffered assay medium (Sigma D5030) with the respective carbon source(s) in absence of CO_2_ for 1 hr. The glycolytic rate was then measured in real time as the extracellular acidification rate (ECAR). After measuring the basal ECAR, 5 μM oligomycin, 2 μM carbonyl cyanide-p-trifluoromethoxyphenylhydrazone (FCCP) and 1 μM rotenone with 1 μM antimycin were injected sequentially.

### Proteomics

N and Rc cells were treated with Hsp90 inhibitors at experimentally determined concentrations that kill 25% of the cells (IC25): gedanamycin (GA) at 500 and 150 nM, PU-H71 (PU) at 800 and 500 nM, pochoxime A (PX) at 5 and 2 μM, respectively, for N and Rc cells. As a control, N and Rc cells were incubated with similar volumes of DMSO. After 24 hrs treatment, cells were harvested by trypsinization and pelleted by centrifugation. Pellets were immediately frozen in liquid nitrogen and shipped in dry ice to the proteomics service.

The iTRAQ experiment was carried out as previously reported [29] at the Beijing Genomics Institute (BGI, Shenzhen, China). Data managing and analysis were also carried out at BGI as previously described [30]. Within an iTRAQ run, differentially expressed proteins were determined based on the ratios and *p* values provided by the software ProteinPilot. ProteinPilot generates *p* values exploiting the peptides used to quantitate the respective protein, and gives a measure of confidence that the ratio is different from 1. Proteins are considered differentially expressed if the ProteinPilot ratios are either > 1.2 or < 0.83 and *p* values are < 0.05. Proteins that met these criteria in at least one of the three biological experiments performed, are listed in S1 File and were used in all further analyses. The mass spectrometry proteomics data have been deposited to ProteomeXchange via the partner repository jPOSTrepo (Japan ProteOme STandard Repository) [31] with the dataset identifier JPST000397 (PXD009055 for ProteomeXchange). The principal component analysis (PCA) of the quantitative proteomic changes of the proteins in common in datasets was generated using the R statistical package Modern Applied Statistics with S (MASS) (R Project; http://www.r-project.org/). The proteomic signatures of Hsp90 inhibition with the different Hsp90 inhibitors were determined using the same R package.

### Network and graph-based analyses

A protein-protein interaction (PPI) network can be represented as a graph where proteins represent nodes (or vertices) and interactions represent edges. We built PPI networks from transformation-enriched programs of Rc cells by integration of PPI data from public databases using a method previously described for Hsp90Int [3]. For this network analysis, we assumed that proteins, which were depleted upon Hsp90 inhibition based on our proteomic data, completely “disappeared” from the PPI network. To measure the impact of this “disappearance”, graph measures were calculated on these networks. The neighborhood connectivity of a node n is the average connectivity of all neighbors of n [32]. The neighborhood connectivity distribution gives the average of the neighborhood connectivity of all nodes n with *k* neighbors for *k* = 0,1,…. Degree distributions, the node degree of a node n is the number of edges linked to n [33]. A node with a high degree is referred to as hub. The node degree distribution gives the number of nodes with degree *k* for *k* = 0,1,…. Each group of data points is fitted to a power law. All topological parameters were computed with the Cytoscape plug-in NetworkAnalyzer [34]. As a control, all the measures were also computed for the impact produced by the removal of five different random lists of proteins with the same size as the Rc-enriched Hsp90-dependent proteome, extracted from a reported full proteome of NIH-3T3 cells [35].

Highly interconnected groups of nodes (subgraphs), which can be considered as protein complexes and functional modules for different processes of interest, were detected in Hsp90-dependent networks using the Cytoscape plugin Mcode [36]. The functional characterization of the Hsp90-dependent networks was performed by mining for overrepresented GO terms associated with the members of the networks using the Cytoscape plugin ClueGO [37].

### Cell death and cell cycle analyses by flow cytometry

The treatments for these analyses were as follows: growth in hypoxia (1% O_2_ environment in a Galaxy 48R incubator from Eppendorf) or low glucose (1 g/l) for 24 hrs; 20 μM AZ628 plus 20 μM PD98059 for 4 hrs; 1 μg/ml cycloheximide for 4 hrs; 1 ng/ml rapamycin for 6 hrs; 10 μM aphidicolin for 15 hrs; 2 mM hydroxyurea for 15 hrs. For some experiments, N cells were transfected with plasmids pEGFP-Q74 or pEGFP-Q23 or grown in medium supplemented with a cocktail of the growth factors insulin (100 ng/ml) and EGF (100 ng/ml) for 24 hrs. After these afore-mentioned pretreatment periods, N and Rc cells were treated with GA (0-1000 nM), PU (0-1000 nM) or PX (0-2000 nM) for an additional 48 hrs. For cell death analyses, cells were harvested and stained with 5 μg/ml propidium iodide (PI) for 15 min at 4°C, and analyzed by flow cytometry. For cell cycle analyses, cells (1 × 10^6^) were harvested and fixed with ice-cold 70% ethanol, washed with phosphate-buffered saline (PBS), treated with 0.1 mg/ml RNAseA in PBS for 5 min, followed by 20 min in PBS containing 5 μg/ml PI. At least 20’000 cells were analyzed with a FACSCalibur flow cytometer and the software CellQuest Pro (BD Biosciences). Percentages of cells in each phase of the cell cycle (G1, S and G2/M) for cell cycle analyses or of PI-positive cells for cell death analyses were evaluated with the software FlowJo (TreeStar). PI-positive cell values are shown as Hsp90 inhibitor dose response curves or as histograms % excess PI-staining (% PI staining with drug - % PI-staining without) on the Y axis.

### Determination of relative protein abundance

Protein abundance of treated and untreated Rc and N cells was determined by measuring protein concentrations with the Bradford assay and with bovine serum albumin as a standard. For each case, triplicate sample of cells were washed with PBS, harvested from culture dishes by trypsinization, counted in a Neubauer chamber; 5 × 10^5^ cells were then pelleted and resuspended in PBS for protein concentration measurement.

### Statistical analyses

Unless indicated otherwise, the data shown are averages from at least three independent experiments and statistical analyses were performed using GraphPad Prism 7 (La Jolla, CA, USA). The differences between the groups were analyzed by two tail Student’s t-test. Error bars represent the standard deviation of the mean and * *p* <0.05 denotes differences between the means of two test groups considered statistically significant.

## Results

### Ras transformation of normal NIH-3T3 fibroblasts

To study genetically comparable and matched normal and cancer cells, we transformed the mouse fibroblast cell line NIH-3T3 (N) by constitutive expression of the *Ha-c-RAS* gene. This yielded the Ras-transformed NIH-3T3 cell line. Note that this model system has been widely described and evaluated [26,38-41]. Immunoblot experiments confirmed the overexpression of Ras protein in four independent clones (S1A Fig). Furthermore, we observed the phosphorylation of key effectors of Ras, the mitogen-activated protein kinases Erk1/2, and the overexpression of caspase-3 in R cell lines, in accordance with the previously observed phenotype of Ras-transformed cells [41]. We chose clone C (Rc) for further studies. As can be seen in S1B Fig, Rc cells display a more acute spindle-like morphology and are smaller when compared to the parental N cells [41]. These Rc cells form foci if they are continuously cultured after reaching confluency, indicating a loss of contact inhibition. As other cancer cells, Rc cells are more energetic than N cells, consistent with a Warburg phenotype (S2 Fig). The extracellular acidification rate (ECAR) profile in complete medium containing glucose, pyruvate, and glutamine showed that Rc cells have an increased glycolytic flux (S2A Fig). When the cells were cultured only with glucose, the basal ECAR was not affected, indicating that the proton production was primarily from glucose in both Rc and N cells (S2B Fig). Addition of oligomycin also showed that Rc cells have a stronger increase and higher maximum glycolytic potential (S2A-B Fig). Overall, these metabolic assays showed that Ras transformation modifies the metabolic profile shifting the normal mouse fibroblasts from a metabolically relatively quiescent to a more energetic state (S2C Fig). When stressed by respiration modulators, N cells compensated with a proportional increase of both glycolysis and oxidative phosphorylation, whereas Rc cells did it with a proportionally more substantial increase in glycolysis to meet the energy demand (compare the slopes of the shift from baseline to stressed activities in S2C Fig).

### Hsp90 inhibitors impact the proteome of both normal and transformed cells

Several earlier studies have taken advantage of the druggability of Hsp90 in order to understand the global changes at the proteome level that the inhibition of this important interactome hub produces in cancer cells [42,15,43-46]. Targeting of Hsp90 has a considerable impact on the structure of cellular protein networks; the interactome is perturbed, becomes laxer and likely less efficient in communicating changes and stimuli to maintain cellular homeostasis [5]. For the purposes of this study, we compared the Hsp90-dependent proteomes of N and Rc cells by mass spectrometry (MS). To minimize off-target and indirect effects of Hsp90 inhibition, we first determined the IC25 concentrations for the three inhibitors and both cell lines (see Materials and methods). For the MS experiment itself, Fig 1A gives a schematic overview of the experimental approach. N and Rc cells were treated with the three chemically very different Hsp90 inhibitors [11] GA [47], PU [48], and PX [49]. Tryptic peptides were labeled with a specific isobaric iTRAQ reagent for multiplexing (Fig 1A and S1 File). The proteomes of N and Rc cells treated with GA showed a similar number of changes, both for protein depletion (Fig 1B) and enrichment (S3 Fig). In terms of numbers of proteins, the outcome was globally similar for the other two inhibitors (Fig 1B). A gene ontology (GO) enrichment analysis highlighted several functions associated with the depleted set of proteins in each cell line (Fig 1B). Comparing N and Rc cells, the set of proteins depleted or enriched in response to treatment with Hsp90 inhibitors is quite different; in fact, for example for GA, they only share 14% and 12% depleted and enriched proteins, respectively (Fig 1C). The differential sensitivity of Rc and N cells to Hsp90 inhibitors, which is reflected in their IC25 values (see Materials and Methods) and will be discussed below, appears not to reside in the number of affected proteins and functions. We next determined how similar the response of Rc cells is qualitatively to the three different inhibitors.

**Fig 1.**
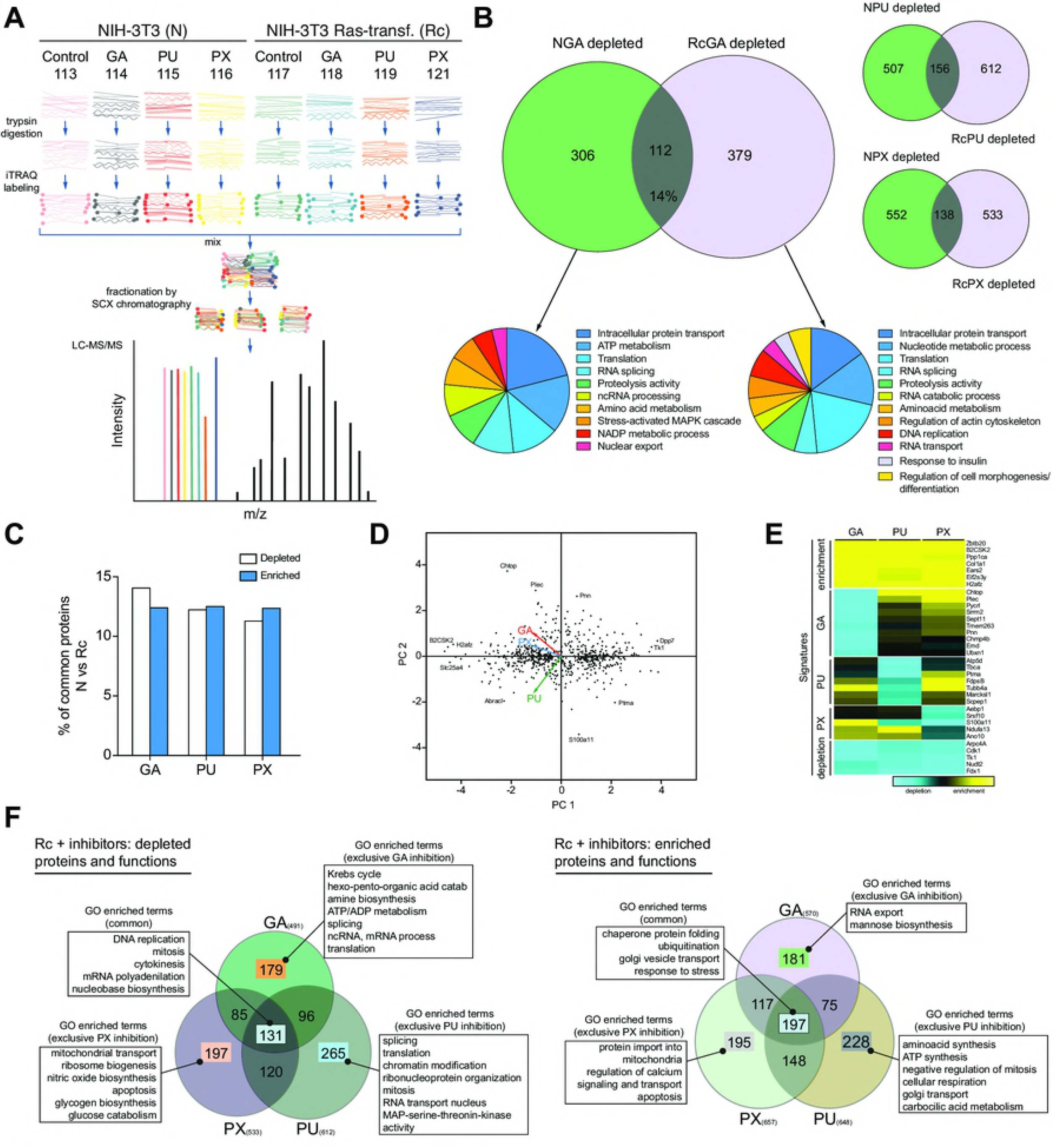
Global analysis of the Hsp90-dependent proteome. (A) Overall experimental design, that shows the steps involved in determining relative quantities of proteins between N and Rc cells, un- or treated with Hsp90 inhibitors, allowing to compare eight samples simultaneously. In LC-MS/MS plot the 8 color peaks express the 8 report ions of iTRAQ. The height peaks show the relative amount of peptides per sample, other black peaks are the MS/MS ion peaks of peptide fragmentations for protein identification. (B) Venn diagrams highlighting the number of proteins depleted in N versus Rc cells treated with three different inhibitors and the GO analysis of those sets of proteins; for GA, the % of proteins in the overlap is indicated, and for all three inhibitors, they are shown in panel C. (C) Protein intersections between N and Rc cells treated with the mentioned Hsp90 inhibitors; the data are from panel B and S3 Fig. (D) Principal component analysis (PCA) of the quantitative proteomic changes of the 656 proteins common to 3 datasets of Rc cells treated with Hsp90 inhibitors. The PCA “loadings” (colored arrows) illustrate the similar/distant behaviors between variables (inhibitors). (E) Proteomic signatures of Hsp90 inhibitors. Heat map depicting the proteins that are up- or downregulated and clustered by common enrichment and depletion signatures for the 3 datasets (top and bottom panels) or distinctive depletion signature for each inhibitor (intermediate panels) on Rc cells. (F) GO functional enrichment analysis of common (or inhibitor-exclusive) affected protein abundance (depletion or enrichment) of Rc treated cells.

Compiling the Log_2_ ratios of the specific isobaric iTRAQ signals from the three datasets coming from Rc cells treated with GA or PU or PX, we could extract a common set of 656 proteins whose abundance was altered by the inhibitors. We used a principal component analysis (PCA) as a dimension-reduction tool to facilitate the evaluation (Fig 1D). The PCA highlights that the cell line Rc responds quite differently to the three compounds; this is apparent from the lack of a large number of proteins at the center of the scatterplot, and the presence of many outliers (proteins) that can define both the extremes of the common response or the specific response towards each inhibitor (Fig 1D). Despite all the differences, one can identify a consistent signature of depleted and enriched proteins that is common to all inhibitors (Fig 1E; very bottom and top of the heat map). In addition, the heat map shows specific signatures for each inhibitor (Fig 1E, central panels). A GO enrichment analysis of the cellular functions associated with depleted and enriched proteins saliently demonstrates how each inhibitor affects different sets of essential functions for cancer survival, despite the fact that they target the same binding pocket of the same protein (Fig 1F).

### Hsp90 inhibitors reshape the proteomic changes associated with oncogenic transformation

The emphasis of the analyses presented above was on comparing cells with and without treatment with Hsp90 inhibitors. Next, we decided to study the changes that take place during the malignant transformation, which is associated with an increased sensitivity of Rc cells to Hsp90 inhibitors. We recently reported a technically similar study with myoblasts differentiating into twitching myocytes in culture; we had found that a vast group of proteins involved in the establishment and maintenance of the skeletal muscle state are severely affected by Hsp90 inhibition with GA [50]. We wanted to investigate the impact that Hsp90 inhibition has on the portion of the proteome involved in the establishment of the Ras-transformed state. Our quantitative proteomic analysis showed that 697 and 630 proteins are enriched and depleted, respectively, upon transforming N into Rc cells (Fig 2A). We could identify relevant Reactome pathways that are associated with this transformation (Fig 2A). As expected, some that are typical of a cancer phenotype are down-regulated (for example, the TCA cycle and respiratory electron transport, and programed cell death), whereas others are enriched (for example, cell cycle, glucose metabolism and signaling pathways) (Fig 2A). The changes in metabolic pathways are experimentally mirrored by the results of our ECAR analysis discussed above (S2A-C Fig). Importantly, as a result of Hsp90 inhibition, a substantial proportion (about 33% at the protein level) of these proteins associated with Ras transformation are considerably counteracted by the three Hsp90 inhibitors, while smaller percentages (for example, 12 % for GA) of these proteins are enriched (Fig 2B). Hypothetically, the combined treatment with GA, PU and PX could affect up to 53% and 60%, respectively, of the enriched and down-regulated proteins associated with Ras transformation (Fig 2B). If one organizes the proteins that are enriched in Rc cells as a protein-protein interaction (PPI) network, one can simulate how this network would be affected by the complete disappearance of proteins upon treatment with GA, PU or PX. The sole targeting of Hsp90 has a huge impact on the Rc-enriched network topology (Fig 2C). The average neighbor connectivity of network members and the node degree distribution of the network are reduced as result of Hsp90 inhibition. The power law line that fits the points of each dataset shifts downwards and to the left, which means that the protein nodes in the network have become less connected (Fig 2C). This phenomenon is specific to Hsp90 inhibition since the elimination of a random but similarly sized list of proteins on the Rc-enriched network does not affect the network topology (Fig 2C, grey dots and lines). Targeting the Hsp90 molecular chaperone system of the proteo-/interactome of a Ras-transformed cell leads to a looser, disconnected and most likely less functional network. By exploring these networks with Cytoscape [51] and its plugin MCODE, we could find highly interconnected regions (complexes) in the Rc-transformation interactome of which some are illustrated in Fig 2D. These clusters contain functional complexes, which might get disrupted because of the inhibitor-induced depletion of one or several components (Fig 2D). Altogether, these proteomic and bioinformatic analyses reveal that Hsp90 provides an essential support for the newly established Ras transformation program.

**Fig 2.**
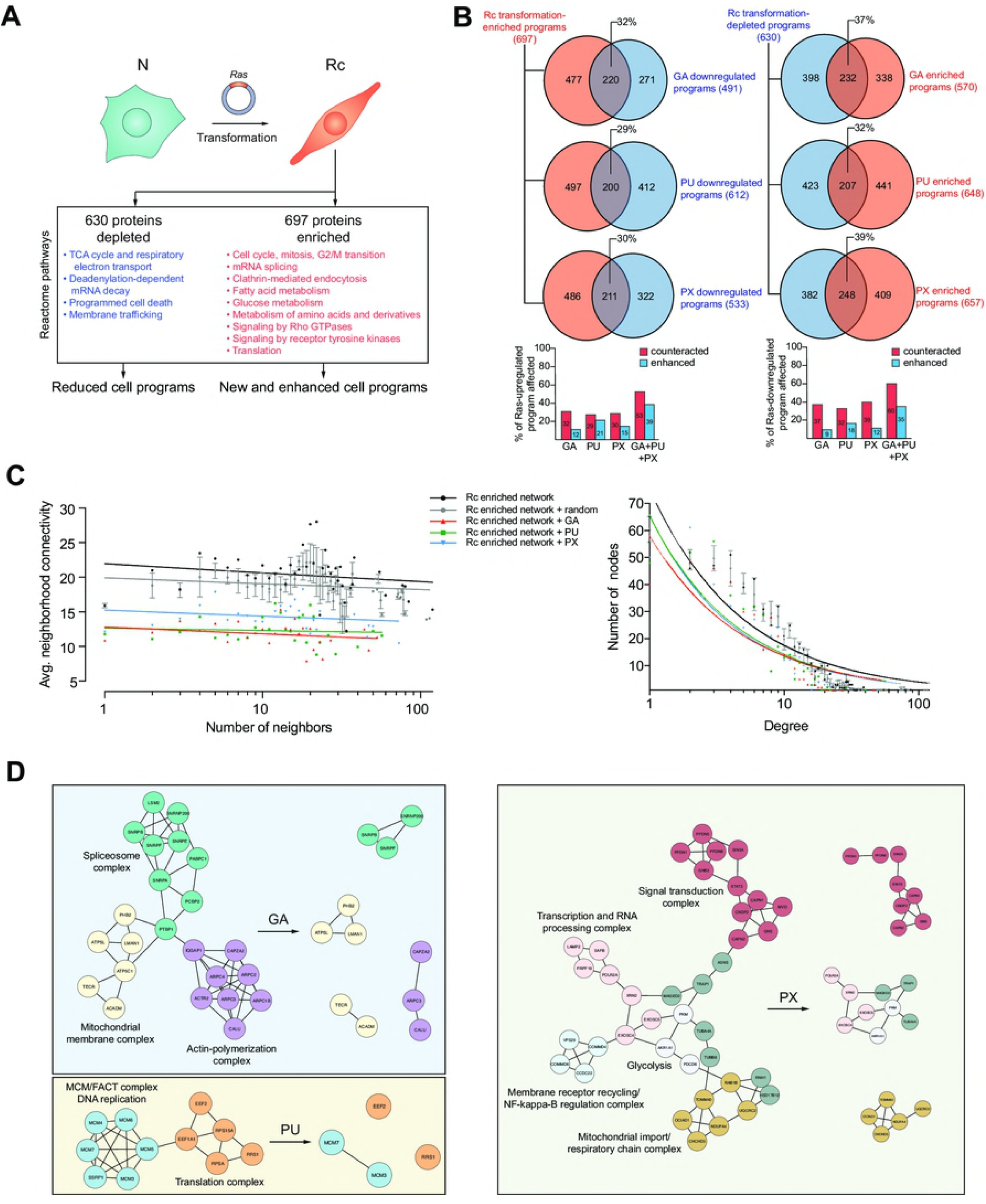
The Hsp90 chaperone system supports and wires the proteome of the transformation program. (A) Scheme of the transformation protocol and of the major proteomic changes; Reactome pathways defined by proteins that are significantly changed in Rc relative to N cells are indicated. (B) Venn diagrams showing the number of proteins of the Rc-specific program that are affected by Hsp90 inhibitors. The colors red and blue indicate upregulated and downregulated proteins. Note that the indicated percentages are calculated by dividing the number of proteins in the intersect by 697 and 630 for the Venn diagrams on the left and on the right, respectively (see panel A), and that in the bar graphs, the values for the combined treatments with all three inhibitors are hypothetical. (C) Topological features of the Rc transformation-enriched network. The effects of the pharmacologically induced depletion of proteins is compared with the effect of removing the same number of proteins, but chosen randomly, in five independent replicates (average effect of five different random lists). Each group of data points is fitted to a power-law. (D) Functional PPI complexes present in Rc cells are disconnected by Hsp90 inhibition. For the purpose of illustrating the impact of Hsp90 inhibition, it is assumed that affected proteins are completely removed.

### The hypersensitivity of Rc cells to Hsp90 inhibitors can be reverted by limiting activated pathways and metabolic resources

Since we had generated the N/Rc pair of cells to study the differential sensitivity to Hsp90 inhibitors, we next ascertained that they indeed display such a differential sensitivity. The general scheme of the experiments is shown in Fig 3A and the inhibitor-induced cell death in Fig 3B. As these dose-response experiments unambiguously demonstrate, Rc cells are substantially more sensitive to the three inhibitors GA, PU, and PX than their parental N cells. This recapitulates and confirms the anecdotal impression that cancer cells are more sensitive than normal cells. We then hypothesized that transformed cells are in some hyperactivated state that confers hypersensitivity to Hsp90 inhibitors. To test this hypothesis, we attempted to revert Rc cells to a more quiescent state by blocking the pathways activated by Ras or by tuning down their metabolism (Fig 3A and C). Specifically, we used a combination of inhibitors, AZ628 as a Raf kinase inhibitor and PD98059 as a MEK inhibitor, targeting two key mediators of the Ras signaling pathway. Rc cell metabolism was manipulated by starvation in low glucose medium or by culturing the cells under hypoxic conditions. The pretreatment of the Rc cells with Ras signaling inhibitors renders Rc cells almost as resistant to GA as N cells (Fig 3C). Low glucose has a similar impact while hypoxia completely eliminates the relative hypersensitivity (Fig 3D and E). To rule out a clonal effect of Rc cells, we repeated the hypoxia experiment with an independent Ras transformant, R cell clone D (Rd); the hypersensitivity of these cells to GA is similarly abolished when they are maintained for 24 hrs in hypoxia and then treated for an additional 48 hrs with GA (S4 Fig). We confirmed the hypersensitivity of Rc cells and the protective effects of hypoxia by examining the increased turnover of a few paradigmatic Hsp90 clients in response to Hsp90 inhibitors by immunoblotting (Fig 3F). Thus, oncogenic transformation does indeed hypersensitize cells to Hsp90 inhibitors, but this can be reverted by manipulating the oncogenic signaling pathway(s) or by reducing metabolic activity.

**Fig 3.**
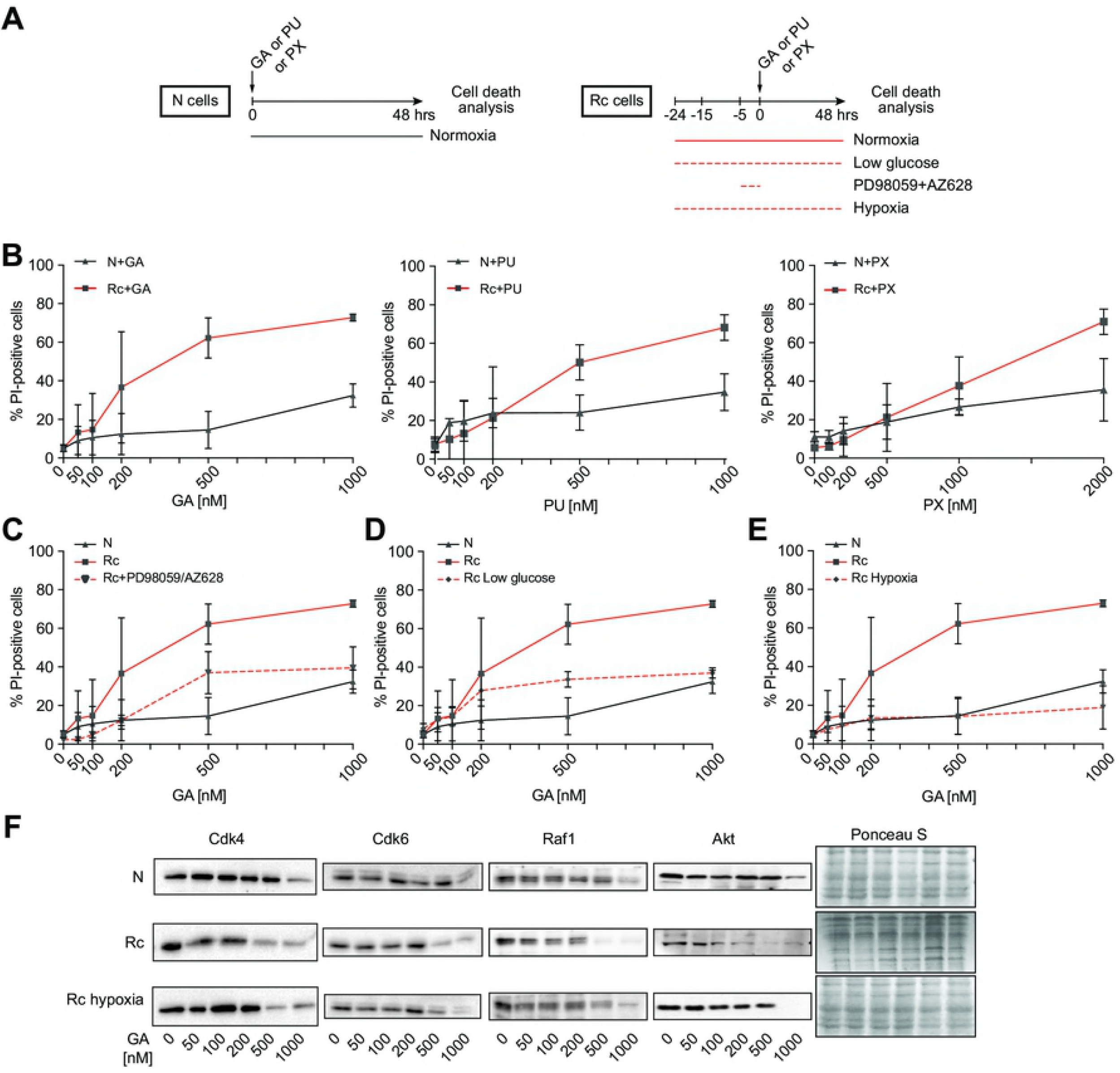
The hypersensitivity of Rc cells to Hsp90 inhibitors can be counteracted by reducing activated pathways and metabolic resources. (A) Scheme of the treatment and analysis regimen. (B) Concentration-dependent induction of cell death of N and Rc cells by GA, PU, and PX after 48 hrs of treatment. (C-E) Impact on GA-induced cell death of Rc pretreated with AZ628 and PD98059 (both 20 μM, 4 hrs) (panel C) or pre-cultured in low glucose (1 g/l) (panel D) or in hypoxia (1% O_2_) for 24 hrs (panel E); low glucose and hypoxic conditions were maintained during the 48 hrs treatment with GA. Note that the data for GA-treated N and Rc cells (solid black and red lines) are identical to those of panel B and reproduced here for better visualization. Averages with standard deviation of 3 experiments are shown. (F) Immunoblots of the indicated Hsp90 clients from cells treated with GA for 24 hrs at the indicated concentrations; the filter stained with Ponceau S serves as loading control.

Considering the dramatic impact of hypoxia on Hsp90 inhibitor sensitivity, we further characterized the effects of hypoxia on Rc cells. It had been reported that hypoxic conditions in the microenvironment of tumors or cells lead to the upregulation of dormancy markers such as Nr2f1 and p27Kip, which in turn induce a growth arrest [52,53]. We found that N cells do express p27Kip and Nr2f1, but that their levels are strongly reduced in Rc cells and reestablished upon exposure of Rc cells to hypoxia for 24 hrs (Fig 4A). Using flow cytometry to quantitate DNA in individual cells, we could confirm that N and hypoxic Rc cells are in a more quiescent state; the proportion of cells in the cell cycle phase G0/G1 compared to cells in the S and G2/M phases is reduced compared to Rc cells (Fig 4B and C). The cell cycle distribution of Rc cells with more than 55% of the cell population in S/G2/M is indicative of a more cycling state (Fig 4C, central bars). These data suggested a connection between a relatively more quiescent state and relative resistance to Hsp90 inhibition.

**Fig 4.**
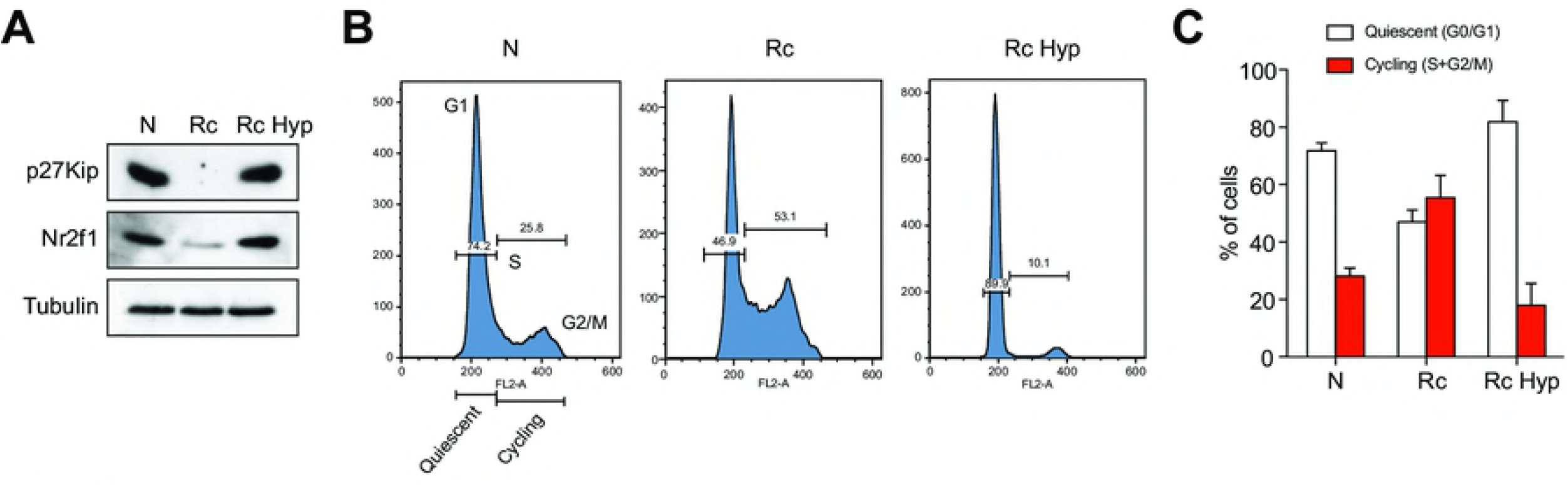
Hypoxia induces a dormant/quiescent state in Rc cells resembling that of N cells. (A) Immunoblots of the dormancy factors Nr2f1 and p27KipN, and tubulin as loading control, from N and Rc cells; Rc Hyp, Rc cells grown under hypoxic conditions for 24 hrs. (B) Flow cytometric DNA analysis of N, Rc and Rc Hyp cells; representative example. (C) Histogram of the cell cycle distribution obtained by flow cytometry. Averages with standard deviation of 4 experiments are shown.

### The balance between relative quiescence and cycling of both N and Rc cells determines their sensitivity to Hsp90 inhibitors

We next asked whether other manipulations of the cell cycle or of proteostasis of both N and Rc cells could influence their sensitivity to Hsp90 inhibition. The specific manipulations and the experimental timing of these experiments are schematically illustrated in Fig 5A. For comparison, hypoxia was included again. Short exposure of cells to low doses of the protein synthesis inhibitor cycloheximide (CHX) can produce a cell cycle arrest [54]. Indeed, the CHX treatment of Rc cells substantially increased the ratio of cells in G0/G1 versus cells in S/G2/M, when tested 4 hrs later (Fig 5B). This increase correlated with a dramatically increased survival of Rc cells exposed to GA for 48 hrs (Fig 5C). Similarly, it has been reported that a short treatment of cells with the inhibitors of DNA replication aphidicolin or hydroxyurea arrest cells in G0/G1 [55]. These drugs provoked the expected result with Rc cells, increasing the quiescent population (Fig 5B) and, at the same time, protecting them from the cytotoxic effects of the GA treatment (Fig 5C). Rapamycin was known to inhibit ribosomal protein synthesis and to lengthen the G1 phase [56]. Rc cells treated with rapamycin also displayed an increase in the fraction of cells in G0/G1 and the accompanying increased survival after Hsp90 inhibition for 48 hrs (Fig 5B and C). Inducing a shift of the relatively rapidly dividing Rc cells to more quiescent cells, more similar to N cells, is sufficient to make these cells less sensitive to Hsp90 inhibition, indeed at similar levels as N cells (Fig 5B and C). Conversely, the stimulation of N cells with a cocktail of the growth factors (GF) EGF and insulin [57], caused an increase in the population of cells in a cycling state (S+G2/M) with a concomitant more than 2-fold increase in cell death upon treatment with GA; the toxicity of GA reached levels similar to that seen with Rc cells (Fig 5D and E). A common hallmark of all of these treatments was that they altered the cellular protein contents. Inhibitors of protein and DNA synthesis significantly decreased the whole cell protein contents of Rc cells (S5A Fig), whereas treatment with GF increased it in N cells (S5B Fig). Overall, we observed a correlation between cellular activity (relative quiescence or cycling) and protein levels with sensitivity to Hsp90 inhibitors.

**Fig 5.**
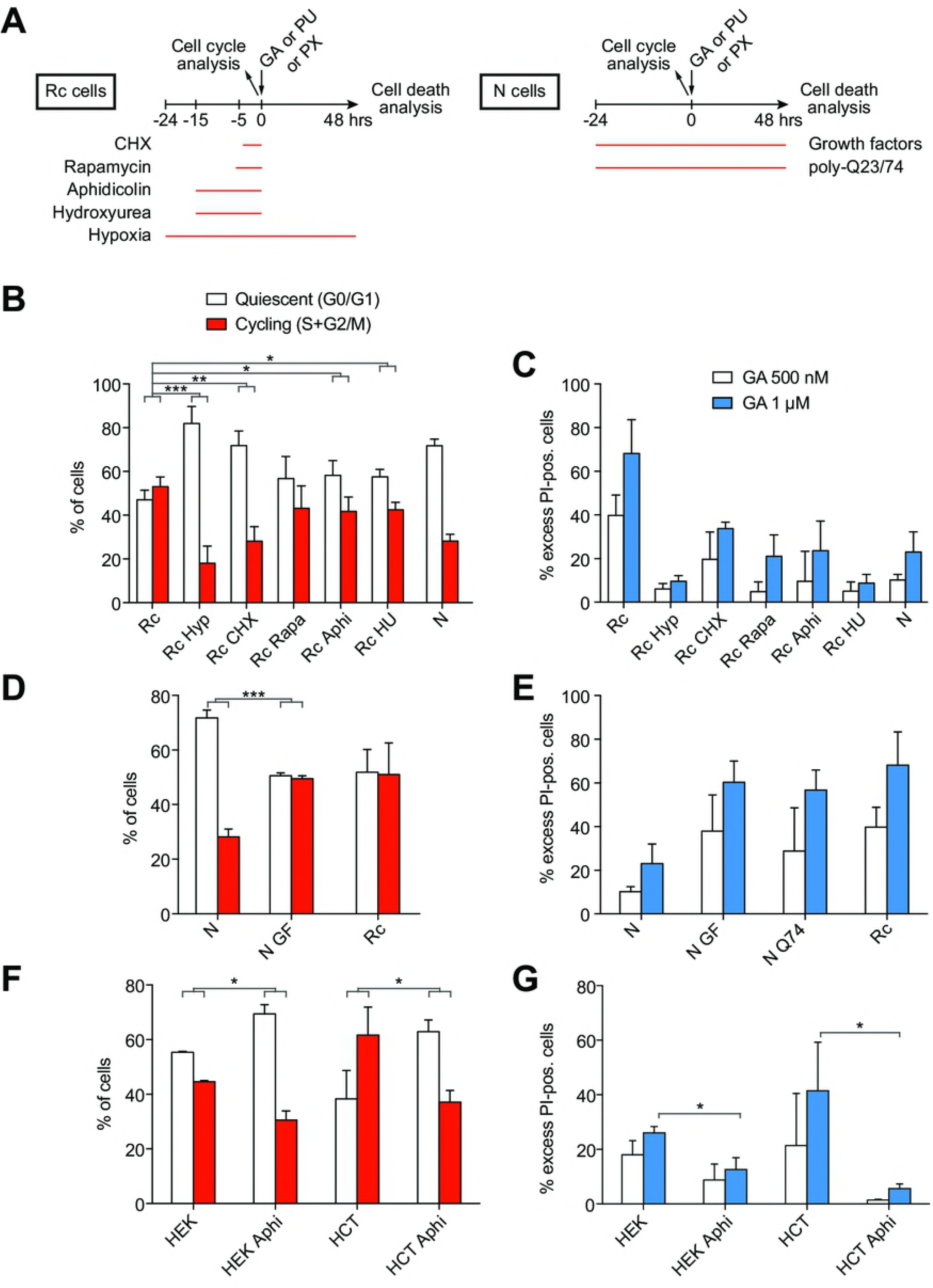
Manipulation of the cell cycle alters the sensitivity to the Hsp90 inhibitor GA. (A) Scheme of the treatment and analysis regimen. (B) Histogram of the cell cycle distribution obtained by flow cytometry of indicated cells subjected to hypoxia (Hyp) or treated with cycloheximide (CHX), rapamycin (Rapa), aphidicolin (Aphi), hydroxyurea (HU); note that the hypoxia data (Rc Hyp) are duplicated here from Fig 4C for comparison. (C) GA-induced death of cells as indicated. (D) Histogram of the cell cycle distribution obtained by flow cytometry of indicated cells; N GF, N cells treated with growth factors. (E) GA-induced death of cells as indicated. N Q74 indicates cells transiently overexpressing EGFP-Q74. (F) Histogram of the cell cycle distribution obtained by flow cytometry of indicated cells; HEK, HEK293T cells; HCT, HCT116 cells. (G) GA-induced death of cells as indicated. Bars represent the means with standard errors of data from three independent experiments. *, significantly different with a *p* value <0.05; ** *p* value <0.05; *** *p* value <0.0005.

Proteotoxic stress is a key feature of established malignant cells [58-60]. Interestingly, by challenging proteostasis of N cells through the expression of a polyQ-expanded and aggregating protein, here an EGFP fusion protein with a stretch of 74 glutamine residues (Q74), we were able to sensitize cells to GA treatment (Fig 5E). A fusion protein with only 23 glutamine residues (Q23), which does not aggregate, served as negative control and did not lead to this sensitization (S6 Fig). Thus, by pushing N cells to cycle more or by perturbing their proteostasis, their relative resistance to Hsp90 inhibitors can be compromised.

### Promoting a more quiescent state in other transformed human cell lines also protects them against Hsp90 inhibitors

To extend this observation to established human cell lines, we treated the human embryonic kidney cell line HEK293T and the human colon cancer cell line HCT116 with the DNA replication inhibitor aphidicolin. This treatment prompted an increase in the population of quiescent (G0/G1) cells and a decrease in the cycling (S+G2/M) ones (Fig 5F). As for the mouse Rc cells, this elicited quiescent states induced a significant relative resistance to GA (Fig 5G). These results strengthen our conclusion that the balance between relative quiescence and cycling determines the relative sensitivity to Hsp90 inhibitors.

## Discussion

The differential sensitivity of cancer over normal cells to Hsp90 inhibitors and thus their differential dependence on Hsp90 has been widely accepted in the community, and yet, the evidence has remained somewhat anecdotal and circumstantial and the mechanisms very poorly understood [12-22]. Nevertheless, this has formed the basis for numerous clinical trials; currently, a search with the keyword “Hsp90” in clinicaltrials.gov yields 124 hits, of which 18 are active and/or recruiting. Here, we have demonstrated with a comparable isogenic pair of normal versus transformed cell lines that the balance between relative cellular quiescence and activity, and not necessarily anything specific to the oncogenic state, is a key determinant of the sensitivity of cells to Hsp90 inhibitors (Fig 6). A wide panel of treatments that affect this balance modify the sensitivity to Hsp90 inhibitors, both of normal and transformed cells.

**Fig 6.**
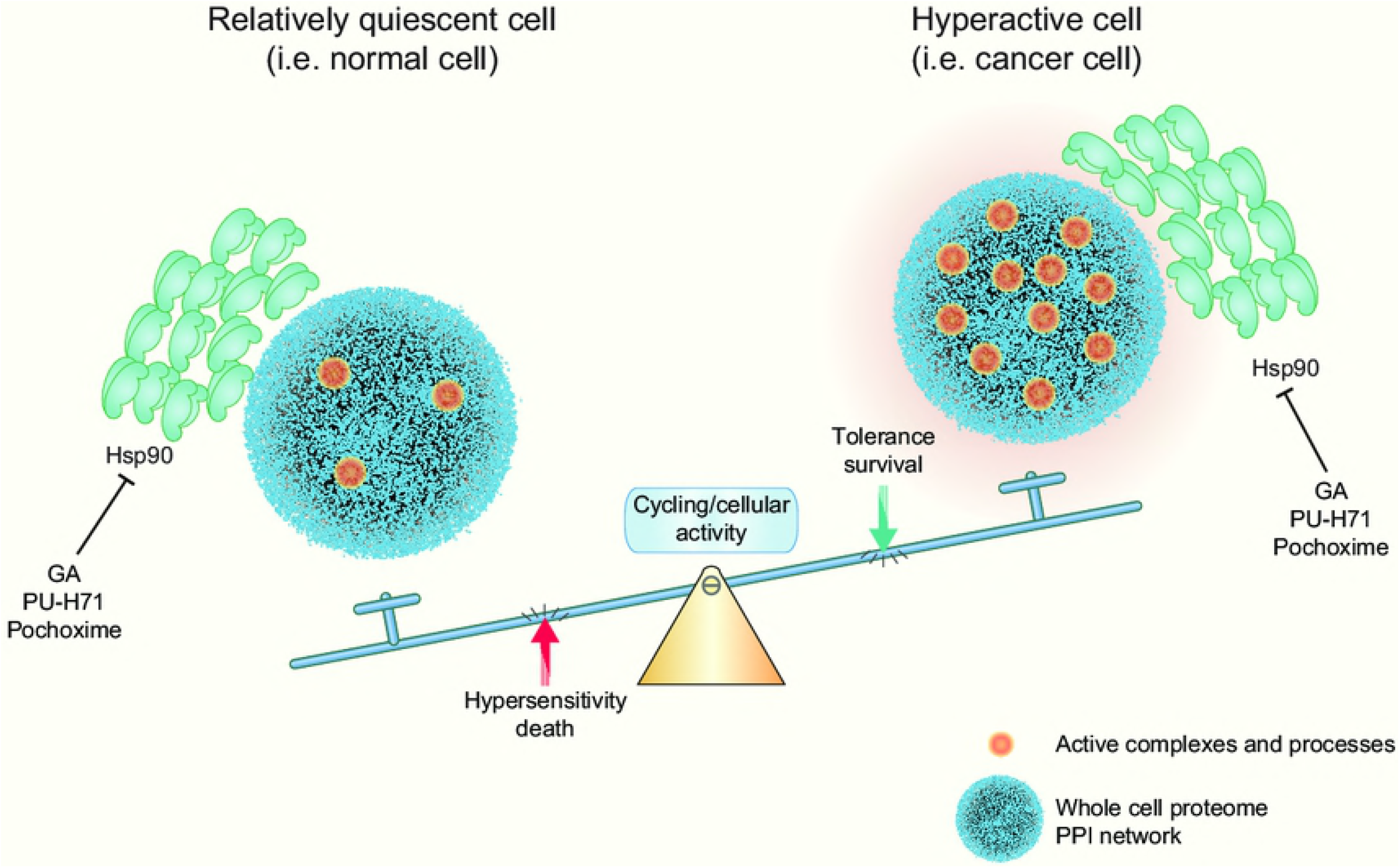
Model of Hsp90 dependence and sensitivity to Hsp90 inhibitors.

The Hsp90 chaperone machine supports many cellular processes that are essential for oncogenic transformation. As a consequence, it has been argued that its capacity to buffer the appearance of polymorphic variants by “chaperoning” variant or mutant proteins, a process which may play an important role in evolution, contributes analogously in oncogenesis [61,6,62-64]. Beyond the support for individual proteins, molecular chaperones in general, and Hsp90 and its cofactors in particular, are essential players to maintain proteostasis. There are numerous illustrations of this role in the literature. For example, the increased sensitivity of cancer cells to Hsp90 inhibitors has also been proposed to be related to their commonly observed aneuploidy and the ensuing proteostatic imbalance [17]. Protein burden, for example imposed by the overexpression of an aggregating protein as in our experiments (Fig 5) or a seemingly innocuous protein as in a recent study in yeast [65], affects cellular fitness and chaperone-dependence. The essential functions of Hsp90 in supporting active cellular states, such as oncogenic transformation, may render cells addicted to it or to direct or indirect regulators of the Hsp90 chaperone machine, and therefore particularly sensitive to Hsp90 inhibitors. For instance, the expression levels of the tyrosine kinase FLT3 in leukemia cell lines is proportionally correlated with the sensitivity to the GA derivative 17-AAG [66]. For acute myeloid leukemia (AML), it was reported that the higher the level of malignancy of these leukemic cells, the higher their sensitivity to Hsp90 inhibition [19]. Moreover, Myc expression, including for AML, was correlated with sensitivity to Hsp90 inhibition [20].

Our global analysis of the Hsp90-dependent proteome showed that for both normal and transformed cells similar numbers of proteins are affected, albeit with relatively limited overlap between the sets of affected proteins (Fig 1B and S3 Fig). Nevertheless, the analysis of the impact of Hsp90 inhibitors on the proteins associated with transformation by the Ras oncogene revealed that a substantial portion of those proteins are affected (Fig 2). Thus, Hsp90 is present to support the basic needs of a normal cell, but when cells are activated or even undergo an oncogenic transformation, the demand for Hsp90 might jump to higher levels. We hypothesize that it is to a large extent this varying degree of Hsp90-dependence or even addiction, which determines the sensitivity to Hsp90 inhibitors.

Our experiments with various treatments affecting the balance between relative cellular quiescence/dormancy and activity demonstrated that both normal and transformed cells can be pushed one way or the other. A particularly interesting and relevant treatment is hypoxia. It is well known that deep and poorly irrigated regions of solid tumors are subject to hypoxia. In primary tumors, hypoxia leads to the appearance of dormant/slow-cycling cells [53]. We found that hypoxia is associated with resistance to Hsp90 inhibitors (Fig 3). Hence, hypoxic areas of solid tumors may be relatively resistant to Hsp90 inhibitors and a potential source for remission when treatment is withdrawn. Primary tumors contain a variety of subpopulations of cells, which differ by their proliferative, cycling and migratory behaviors [67]. It would be interesting to investigate whether and how these different subpopulations are affected by Hsp90 inhibitors. In the context of our findings, cancer stem cells are a particularly intriguing subpopulation of cancer cells [68,69]. Usually quiescent by nature [70], they tend to be relatively resistant to drugs [71,72]. One would expect this resistance to be true for Hsp90 inhibitors as well, although several reports have mentioned that a variety of cancer stems cells can be targeted with Hsp90 inhibitors [73-75]. Since “resistance” is always relative, focused studies are needed to determine whether cancer stem cells are relatively more resistant to Hsp90 inhibitors than the corresponding cycling cancer cells and whether their sensitivity correlates with activity. Intriguingly, it has been proposed that in order to enhance the efficacy of anticancer drugs, it might be necessary to push dormant/quiescent cancer cells, including cancer stem cells, back into the cell cycle, for example with the use of an inhibitor of the protein phosphatase 2A [76]. The risks of reactivating cancer stem cells will need to be carefully balanced against the gain in efficacy of a particular anticancer drug [77]. In any case, if Hsp90 inhibitors are ever to make it to the clinic, these would be important parameters to understand.

## Acknowledgements

The Seahorse X-24 instrument was purchased with the R’Equip grant 316030_145001 of the Swiss National Science Foundation (SNF). In addition, this work was supported by a grant from the SNF to DP, and the Canton de Genève.

## Supporting information

**S1 Fig. Characterization of Ras-transformed cells.** (A) Immunoblot analysis of four independent clones of Ras-transformed NIH-3T3 cells. The panel at the top shows the Ponceau S-stained nitrocellulose filter and indicates equal protein loading. Note that all other presented experiments, except for those of S4 Fig, were done with clone C (Rc cells). Casp3, caspase-3. (B) Phase contrast micrographs of the parental N and the transformed Rc cells.

**S2 Fig. Comparison of energy metabolism of N and Rc cells.** (A) ECAR profiles of N and Rc cells assayed in complete medium containing glucose, pyruvate and glutamine, and treated sequentially with the indicated metabolic inhibitors. (B) ECAR profile of cells assayed in medium with only glucose. The graphs represent the means ± SEM (n = 3). (C) Cell energy profiles with ECAR (a measure of glycolysis) on the X-axis and the oxygen consumption rate (OCR; a measure of mitochondrial respiration) on the Y-axis. The extremes of the four quadrants define the extremes of the different energetic states. The “stressed phenotypes” are the ones in the presence of the metabolic inhibitors; the values used for ECAR and OCR are the highest ones after the injections of oligomycin and FCCP, respectively.

**S3 Fig. Venn diagrams highlighting the number of proteins enriched in N versus Rc cells treated with the three different inhibitors.** The % of proteins in the overlap is indicated, and displayed as a histogram in Fig 1C.

**S4 Fig. GA sensitivity of an independent R cell clone in normoxia and hypoxia.** The graph of this cell death analysis with R cell clone D shows averages of two experiments.

**S5 Fig. Altered protein contents as a function of treatment.** (A) Impact of treatments on protein contents of Rc cells. (B) Impact of growth factors on N cells.

**S6 Fig. Impact of challenging proteostasis with aggregating proteins.** Cell death analysis of N cells transfected with pEGFP-Q23 or pEGFP-Q74, 24 hrs before GA treatment for 48 hrs.

**S1 File. Excel file with the proteomics data of the differentially expressed proteins.** The criteria are those mentioned in Materials and methods.

